## Supplementary Figs for "Uncoupling the roles of firing rates and spike bursts in shaping the STN-GPe beta band oscillations"

### Supporting information

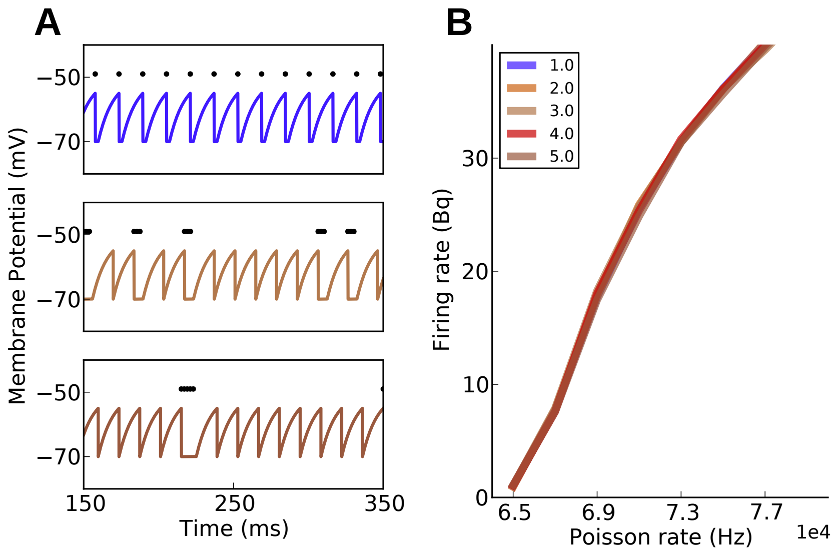


**Fig S1.** State dependent Stochastic Bursting Neuron (SSBN) model **(A)** Membrane potential and spiking pattern for different number of spikes per burst.**(B)** Input current and output firing rate $(f - I)$ curve of the SSBN for different number of spikes per burst.

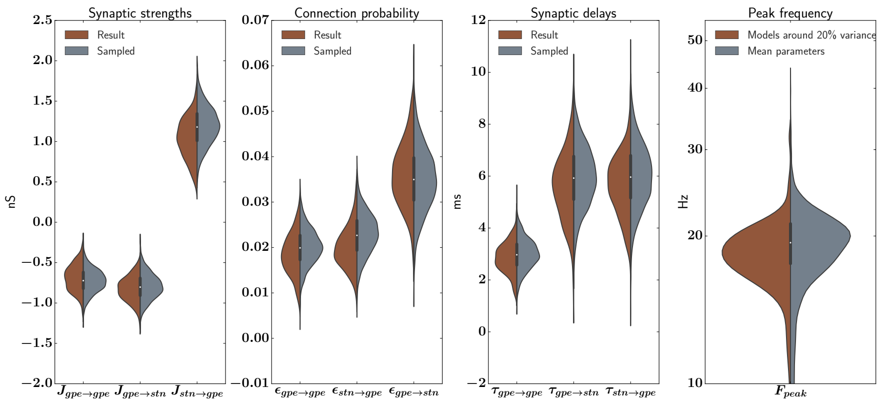


**Fig S2. Robustness analysis.** The areas in gray color of the violin plot shows the distribution that was sampled for robustness analysis. The areas in brown color of the violin plots show the distribution of the parameters that qualitatively reproduce the key results shown in the Fig 3A. See Methods for more detail.

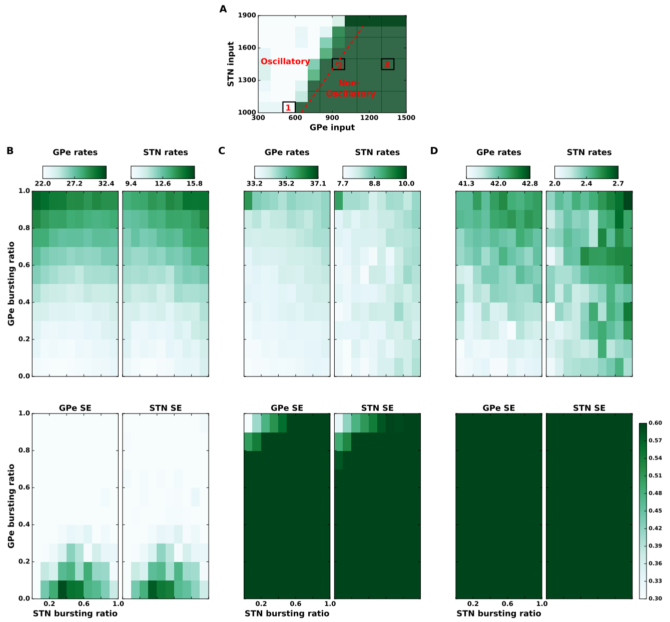


**Fig S3. Effect of spike bursting on STN-GPe network oscillations for an example network from robustness analysis. (A)** Spectral entropy as a function of input to the STN and GPe neurons for a different set of model parameters than used in Fig 3. In this panel the location of the red dotted line and the three exemplary activity regimes are marked in the same place as in Fig 3A for the ease of comparison. **(B):Top** Same as Fig 3B-top. GPe (left) and STN (right) firing rates as a function of the fraction of spike bursting neurons in the STN (x-axis) and GPe (y-axis), in the regime 1. **(B):Bottom** Same as Fig 3B-bottom in the main text. However note that, in this network, regime 1 is on the border and hence shows the non-monotonic effect of STN spike bursting on oscillations as observed in Fig 3C-bottom. **(C)** Same as Fig 3C in the main text. However note that in this network regime 2 is deeper into non-oscillatory regime as compared to Same as Fig 3C. Hence, the effect of spike bursting on oscillations is close to being ineffective. **(D)** Same as in the panel Fig 3D in the main text. These results are qualitatively similar to the ones shown in the Fig 3. Here, however we have used a different set of parameters than the Fig 3 ($J_{gpe-gpe}$ = -0.67, $J_{gpe-stn}$ = -1.0, $J_{stn-gpe}$ = 1.04, $\epsilon_{gpe-gpe}$ = 0.02, $\epsilon_{stn-gpe}$ = 0.02, $\epsilon_{gpe-stn}$ = 0.03, $\tau_{stn-gpe} = 5.96ms$, $\tau_{gpe-gpe} = 3.14ms$, $\tau_{gpe-stn} = 5.34ms$.

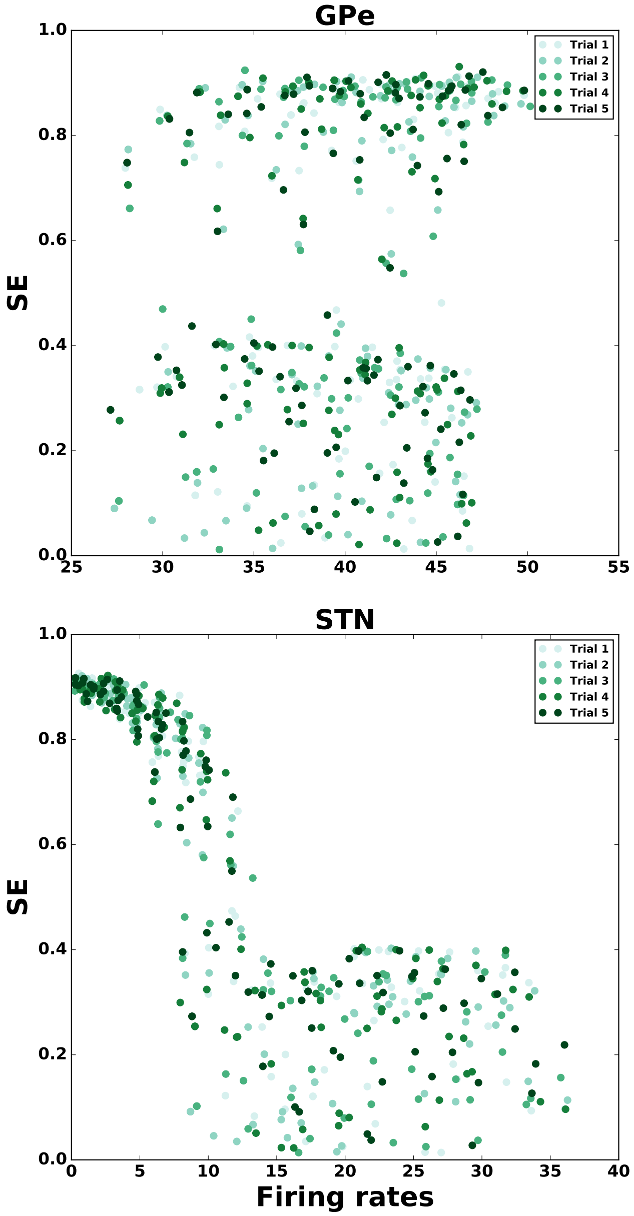


**Fig S4.** Spectral entropy as function of GPe (Top) and STN (Bottom) firing rates. Different colors indicate five different trials with same parameters. Different dots correspond to network simulations with different parameters. For values of GPe firing rates the network can be in an oscillatory or non-oscillatory states, however, high firing rate in STN is necessary to induce oscillations.

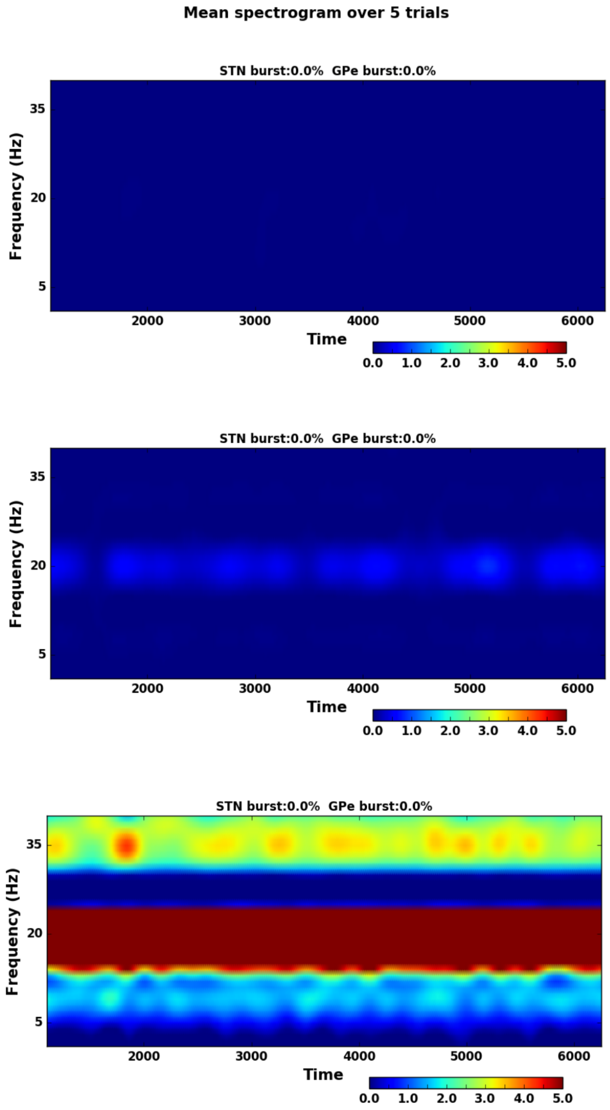


**Fig S5. Spectrograms of network activity in three exemplary network activity regimes. top:** Non-oscillatory regime (marked as 3 in Fig 3A). **middle:** Transition regime (marked as 2 in Fig 3A). **bottom:** Oscillatory regime (marked as 1 in Fig 3A in the main text).

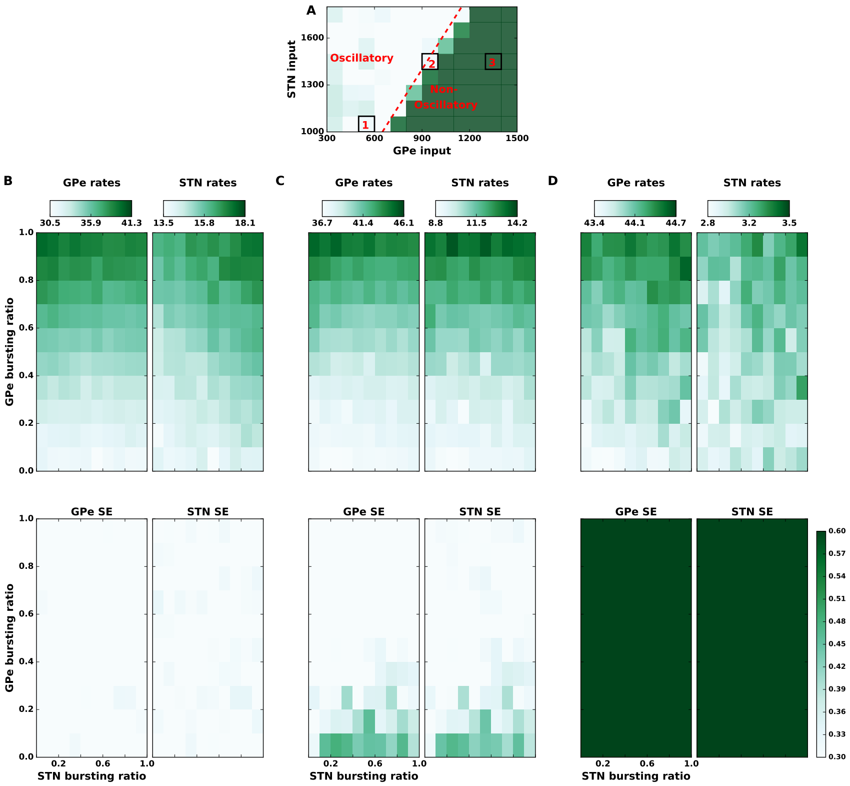


**Fig S6. Reproduction of results shown in Fig 3 for a smaller intra-burst inter-spike-interval ($B_{isi} = 3ms$).** The positions of the regimes 1, 2 and 3 as well as the dashed line dividing the oscillatory and non-oscillatory regime are kept same as in Fig 3A in the main text. Decreasing the within burst inter-spike-interval resulted in reduction in the area of non-oscillatory regime.

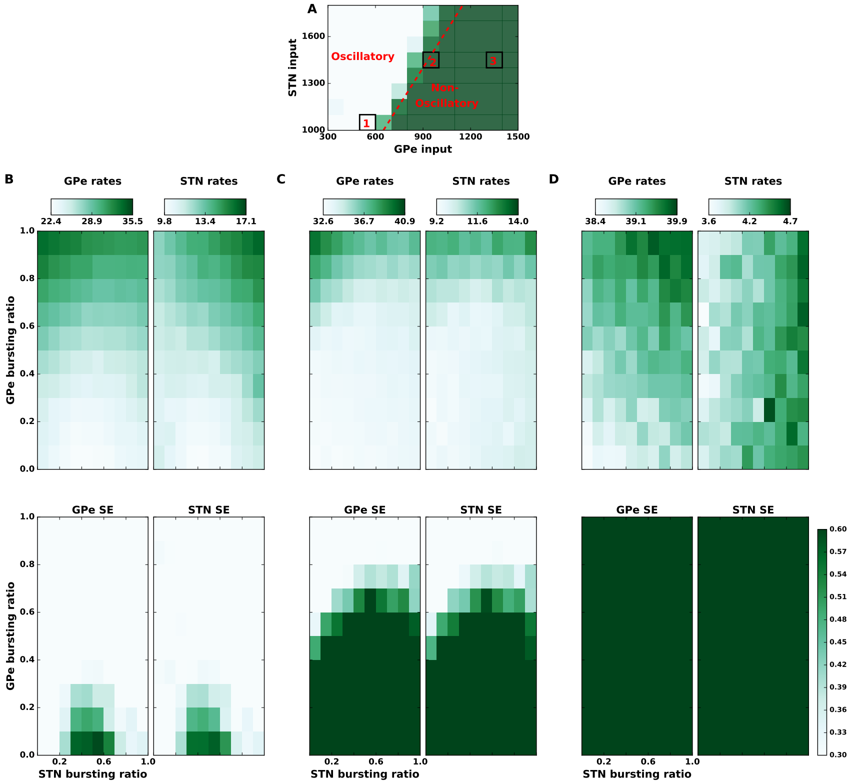


**Fig S7. Reproduction of results shown in Fig 3 for a larger intra-burst inter-spike-interval ($B_{isi} = 7\,ms$.)** The positions of the regimes 1, 2 and 3 as well as the dashed line dividing the oscillatory and non-oscillatory regime are kept same as in Fig 3A. Increasing the within burst inter-spike-interval reduced the region of the oscillatory regime.

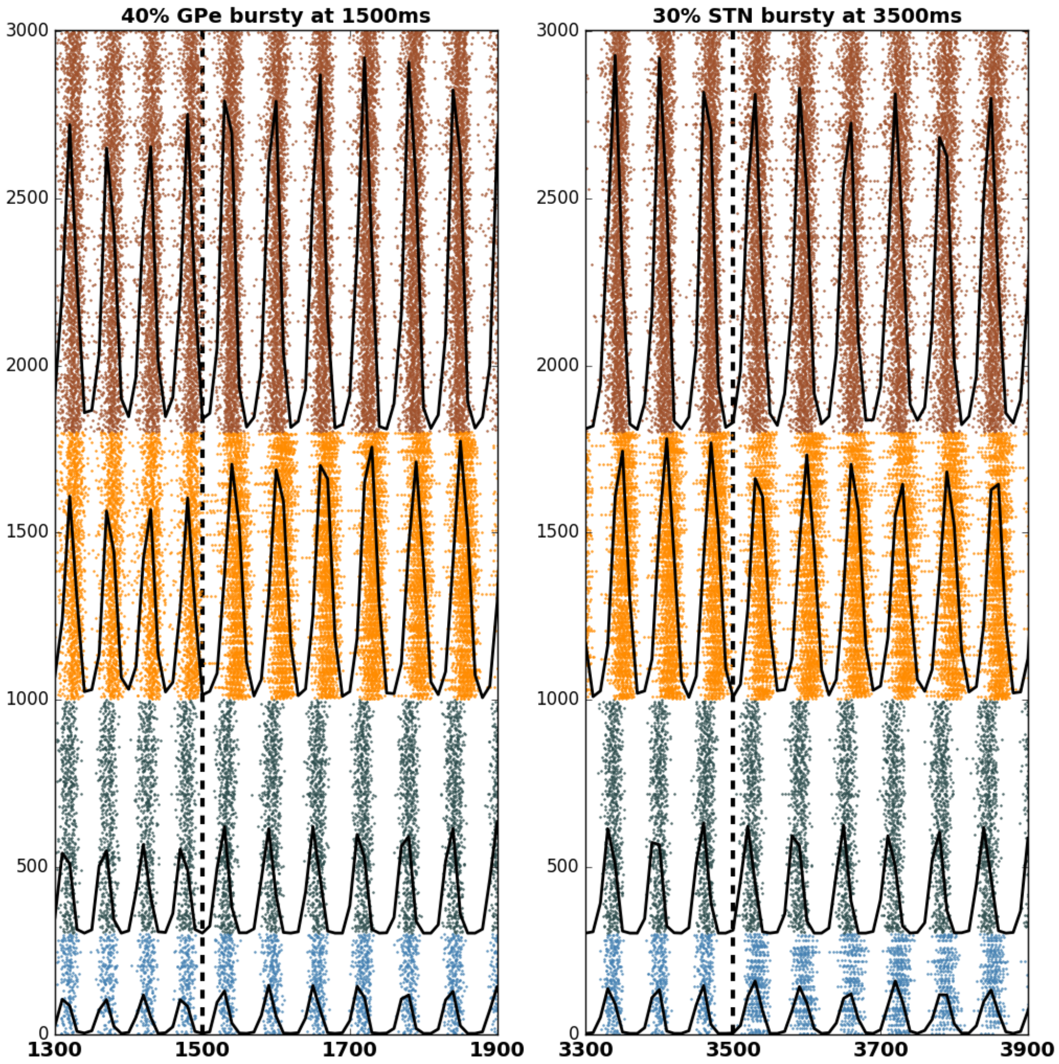


**Fig S8.** Effect of spike bursting when the network was operating in an oscillatory state (regime 1). 40% of GPe neurons (golden yellow) were converted into bursting neurons at time 1500ms - this had no effect of on the network activity state. To see the effect of spike bursting in STN neurons, in addition to the 40% GPe neurons, we also converted 30% of STN neurons (cyan) in bursting neurons at 3500ms. Even this change failed to alter the network activity state. The instantaneous firing rate (binsize = 10 ms) is plotted in black for bursting and non-bursty populations for GPe and STN.

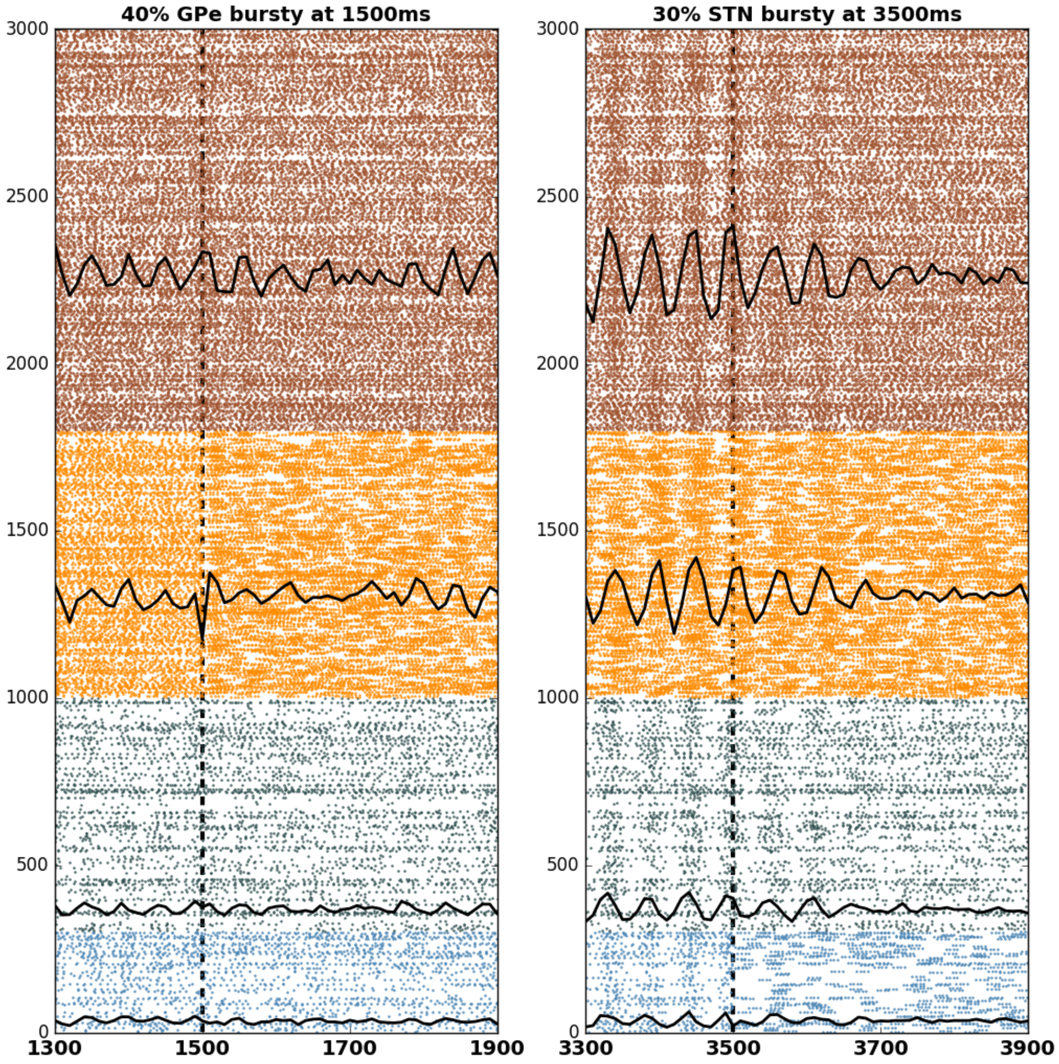


**Fig S9.** Effect of spike bursting when the network was operating in the transition regime (regime 2). 40% of GPe neurons (golden yellow) were converted into spike bursting neurons at time 1500ms. This led to the emergence of weak beta band oscillations (see the spike raster in the right panel before 3500ms). To see the effect of bursting in STN neurons, in addition to the 40% GPe neurons, we also converted 30% of STN neurons (cyan) in bursting neurons at 3500ms. Spike bursting in STN quenched the oscillation initiated by spike bursting in the GPe. The instantaneous firing rate (binsize = 10 ms) is plotted in black for bursting and non-bursty populations for GPe and STN.

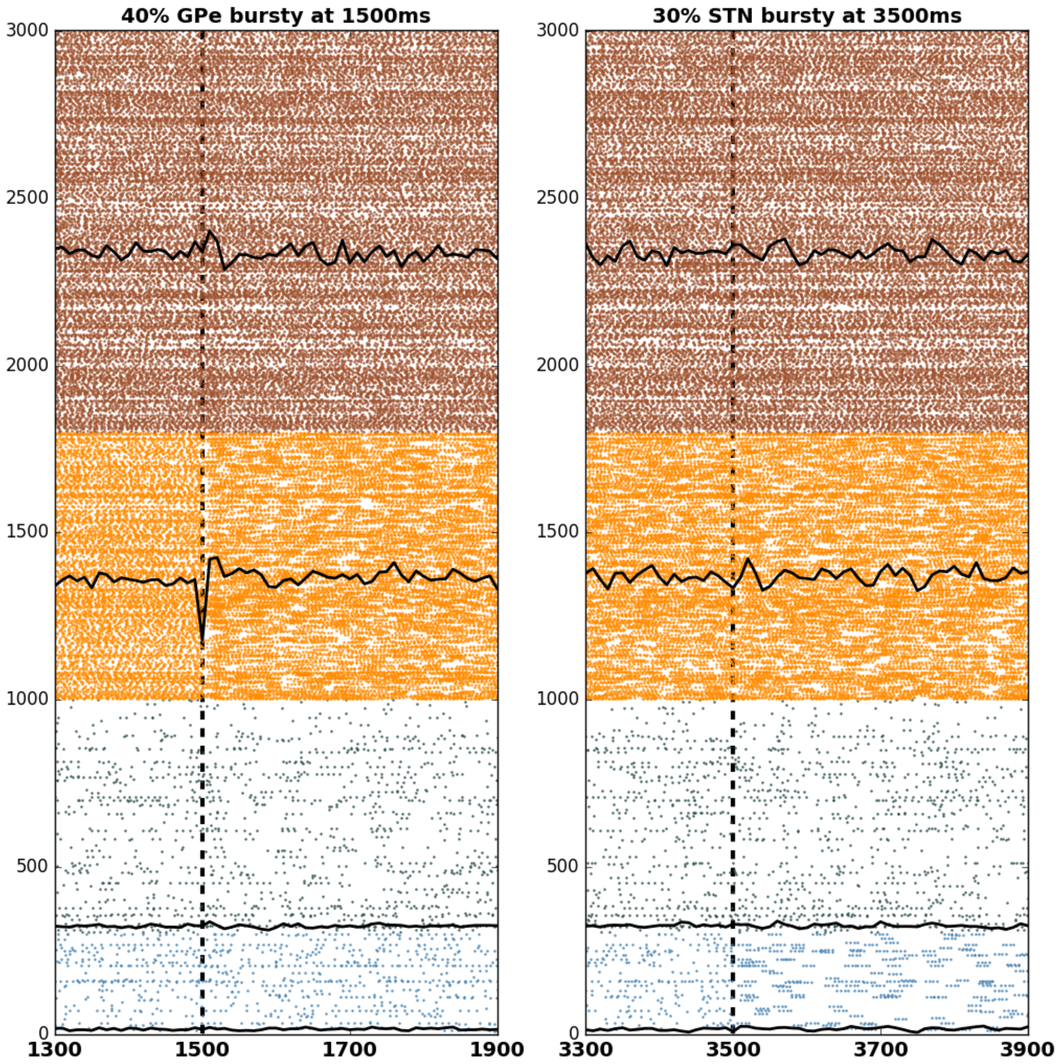


**Fig S10.** Effect of spike bursting when the network was operating in an non-oscillatory state (regime 3). 40% of GPe neurons (golden yellow) were converted into bursting neurons at time 1500ms - this had no effect on the network activity state. To see the effect of spike bursting in STN neurons, in addition to the 40% GPe neurons, we also converted 30% of STN neurons (cyan) in bursting neurons at 3500ms. Even this change failed to alter the network activity state. The instantaneous firing rate (binsize = 10 ms) is plotted in black for bursting and non-bursty populations for GPe and STN.

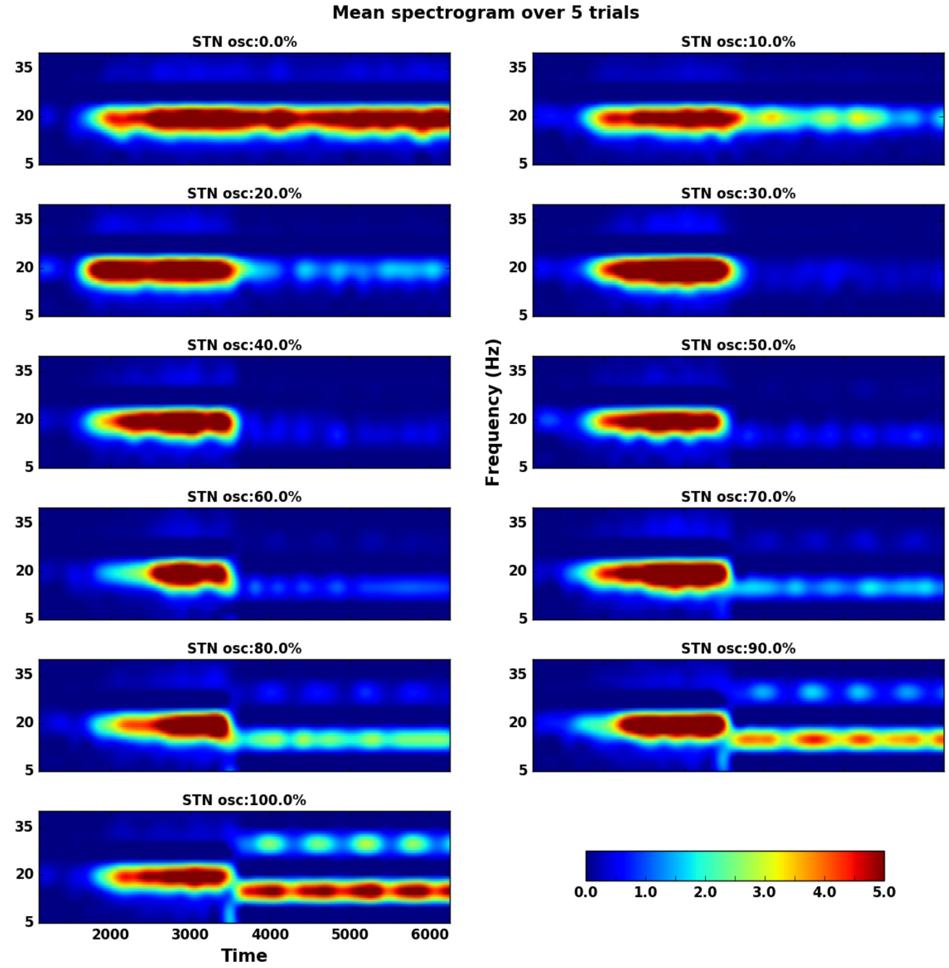


**Fig S11.** STN spike bursting quenches oscillations by imposing a lower frequency on STN population. At 3500 ms, an oscillation of 15Hz was imposed on STN population, instead of replacing STN neurons by bursting neurons. These change in the beta band oscillations because of the injection of 15 Hz oscillations in a fraction of STN neurons are qualitatively similar to the results show in Figure 4. These results explain how spike bursting in STN can quench oscillations when a small fraction of neurons are bursting.

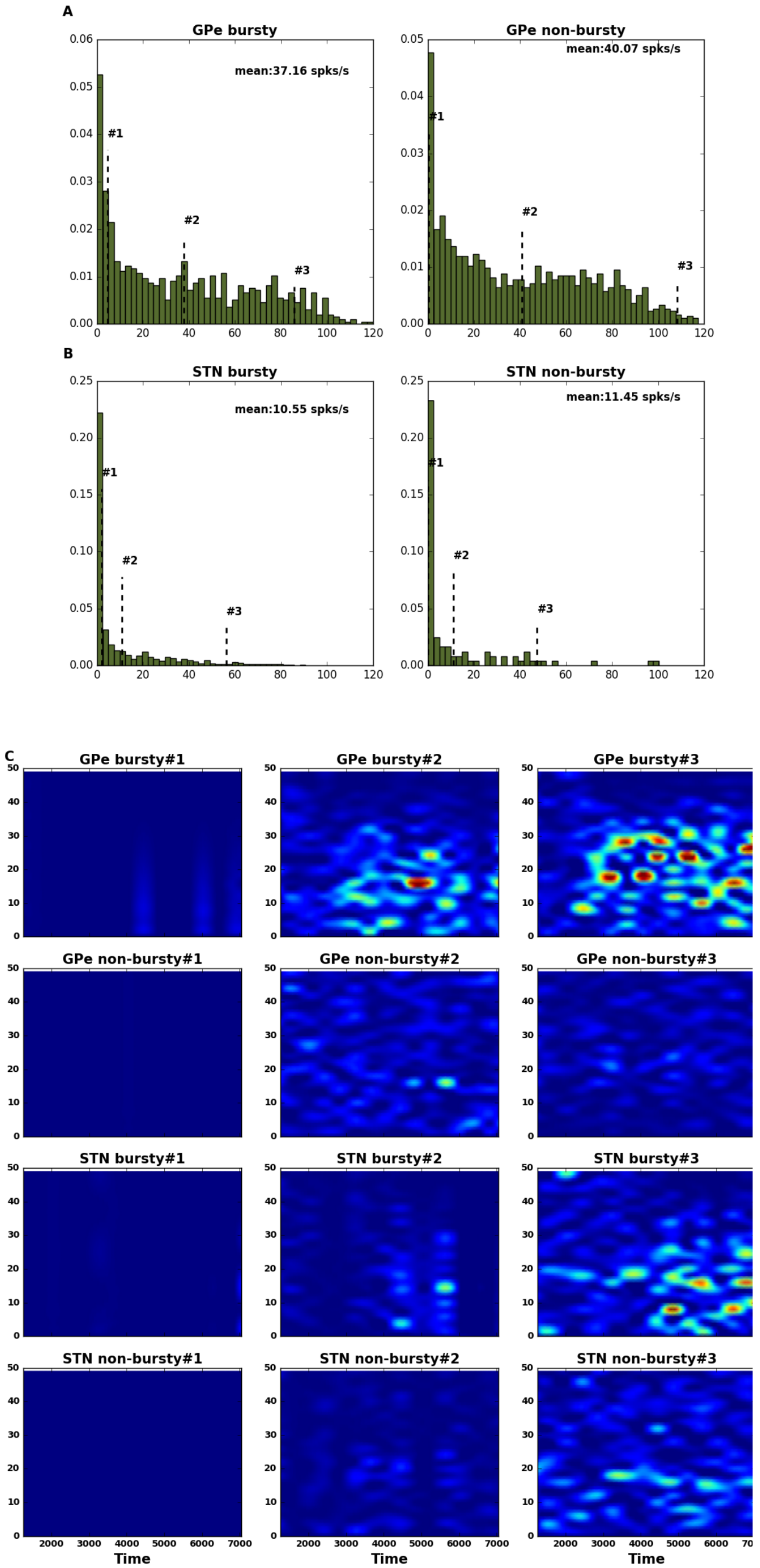
A
GPe bursty
mean:37.16 spks/s
#1
#2
#3
GPe non-bursty
mean:40.07 spks/s
B
STN bursty
mean:10.55 spks/s
STN non-bursty
mean:11.45 spks/s
C
GPe bursty#1
GPe bursty#2
GPe bursty#3
GPe non-bursty#1
GPe non-bursty#2
GPe non-bursty#3
STN bursty#1
STN bursty#2
STN bursty#3
STN non-bursty#1
STN non-bursty#2
STN non-bursty#3
Time

**Fig S12**

**Spectrograms for single neurons with 40% of bursty neurons in GPe and 90% of bursty neurons in STN. (A)** Firing rate histograms of bursty (left) and non-bursty (right) neurons in the GPe. **(B)** Firing rate histogram of bursty (left) and non-bursty (right) neurons in the STN. For both GPe and STN we chose three exemplary neurons, #1 - neuron with average firing rates $\leq$ mean population firing rate (37.16 spks/s), #2 - neuron with average firing rate = mean population firing rate, #3 - neuron with average firing rate > mean population firing rate. **C**: Spectrograms of each of the six chosen neurons from the STN and GPe.
